# Integrated 16S rRNA Bioinformatics Analysis of Gut Microbiota, Predicted Pathways, and Candidate Biomarkers in Inflammatory Bowel Disease

**DOI:** 10.64898/2026.10.16.759573

**Authors:** Aman Zehra, Elamathi Natarajan

## Abstract

Inflammatory bowel disease (IBD) is associated with changes in gut microbiota composition and predicted microbial functions that may relate to intestinal immune processes. This study compared gut microbiota profiles between individuals with IBD and healthy controls and examined predicted microbial pathways and their associations with bacterial genera.

Publicly available 16S rRNA amplicon sequencing data from 22 stool samples (11 IBD and 11 healthy controls) were analysed. Taxonomic profiling, diversity analysis, differential abundance testing, and PICRUSt2-based functional prediction were performed. Microbe–pathway associations were examined using Spearman correlation and Cytoscape. Extended analyses integrated differential abundance with machine-learning feature importance to prioritize candidate genera, evaluated a three-genus signature, assessed selected findings in an independent dataset, and compared predicted fermentation- and acetate-related pathways.

Alpha diversity was significantly lower in the IBD group (Shannon index, p = 0.003), and Bray–Curtis PCoA showed separation of the two groups mainly along PC1 (PERMANOVA pseudo-F = 6.125, p < 0.001), although this separation may partly reflect differences between the source cohorts. DESeq2 identified 44 significantly differentially abundant genera among 186 tested (FDR-adjusted p < 0.05), of which 32 were lower and 12 were higher in the IBD group relative to healthy controls; 27 of these genera were detected in only one of the two groups. PICRUSt2 predicted lipopolysaccharide and peptidoglycan biosynthesis pathways; after normalising pathway abundances for sequencing depth, Spearman analysis identified 84 significant microbe–pathway associations, 47 with lipopolysaccharide biosynthesis (44 positive; ρ up to 0.90) and 37 with peptidoglycan biosynthesis (33 negative; ρ down to −0.82). Five genera were prioritized by integrated biomarker analysis. The combined three-genus signature achieved an ROC–AUC of 0.959. External validation was limited to Bifidobacterium, whose abundance direction was concordant with the discovery analysis. Of the two SCFA-related pathways examined, the predicted acetate-related pathway (methanogenesis from acetate) was lower in the IBD group (FDR = 0.002), whereas the predicted fermentation pathway did not differ significantly between groups (FDR = 0.076).

These findings provide an integrated view of microbial composition and predicted functional potential in IBD. The small sample size, differences in cohort and sequencing characteristics between the IBD and control samples, limited external validation, and reliance on predicted rather than directly measured functions mean that the results are exploratory and require confirmation in larger independent cohorts and through direct functional or metabolomic analyses.

## 1 Introduction

Inflammatory bowel disease (IBD) is classified into two major categories, mainly Crohn’s disease (CD) and ulcerative colitis (UC), and both are referred to as chronic inflammatory disorders of the gastrointestinal tract [1]. The progression of IBD includes several factors like environmental factors, immune responses, combined influence of genetic susceptibility and the intestinal microbiota [1]. Growing evidence suggests that IBD is closely linked to gut microbiome, with alterations in gut microbiome composition and functional characteristics being associated with disease development [2]. The association between IBD and the gut microbiome is complex and bidirectional, reflecting that alterations in microbial communities may affect disease-related processes [2].

The gut microbiota plays a significant role in maintaining intestinal homeostasis [3]. Gut microbial diversity and community composition may change in individuals with IBD. Such alterations in the gut microbial community are often described as dysbiosis, and a reduction in microbial diversity has been widely reported in IBD [3,4]. Several studies have shown changes in the composition of the gut microbiome in individuals with IBD compared with healthy individuals, and changes in the abundance of particular bacterial groups have also been observed [3,4]. However, the relationship between dysbiosis and IBD is complex, and dysbiosis should not be considered a definitive cause of IBD [3].

The relationship between the gut microbiota and the intestinal immune system plays an important role in maintaining intestinal immune homeostasis [2,5]. This relationship is complex because the microbial community can influence immune responses, while the immune system can also shape the gut microbial community [2,5]. In IBD, dysregulation of these interactions may contribute to abnormal immune responses and intestinal inflammation [5]. Hence, understanding the relationship between the gut microbiota and the host immune system may provide insights into the mechanisms underlying IBD pathogenesis [2,5].

Sequencing-based techniques are widely used to characterize gut microbial communities. Among these approaches, 16S rRNA gene amplicon sequencing is frequently used to study bacterial community composition, taxonomy, and diversity [6]. However, 16S rRNA sequencing primarily provides taxonomic information and does not directly measure microbial functional activity [7]. Therefore, PICRUSt2 can be applied to 16S rRNA marker-gene data to predict the functional potential of the microbial community [7].

Alterations in the gut microbiome in IBD are not only limited to changes in microbial composition but also include changes in microbial functions [2,4]. The functional properties of gut microorganisms can affect many biological processes within the intestinal environment [4]. Thus, functional profiling provides additional information beyond taxonomic composition and may assist in identifying microbial pathways potentially associated with disease-related biological processes [2,4,7]. Some microbial pathways are associated with host-microbe interactions because bacterial components, such as lipopolysaccharide (LPS) and peptidoglycan, can interact with the intestinal immune system [5,8,9]. Host pattern-recognition receptors (PRRs) recognize these bacterial components and can influence the immune response [9]. Therefore, analyzing microbial pathways involved in the biosynthesis of these bacterial components may provide insights into potential microbiota–immune interactions in IBD (Figure 1) [5,8,9]. The proposed relationship between IBD-associated microbial dysbiosis, predicted microbial pathways, and potential host immune responses is summarized in Figure 1.

**Figure 1.**
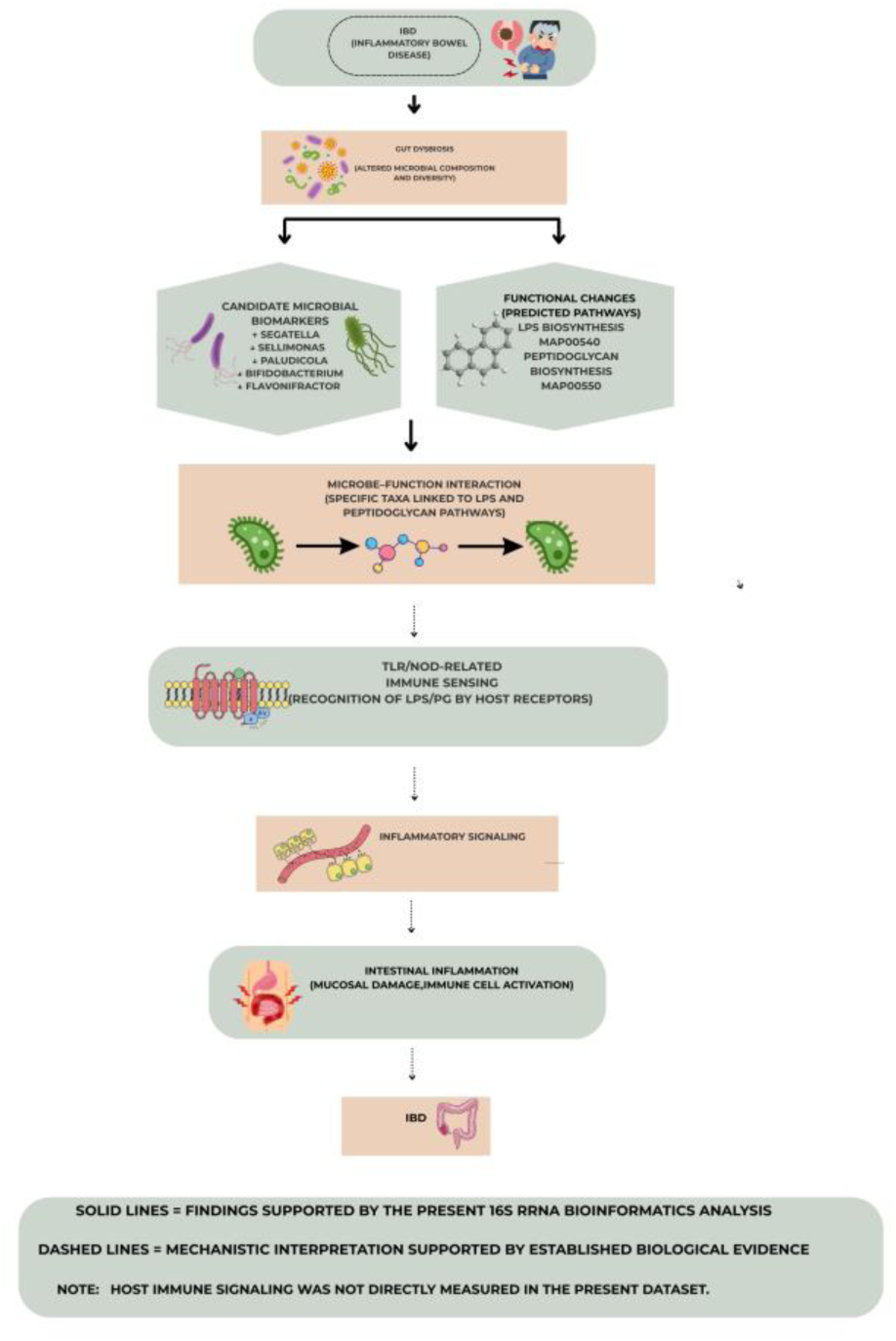
Conceptual overview of IBD-associated gut microbial dysbiosis, candidate biomarkers, predicted microbial pathways, and potential host-microbiota immune interactions. Solid arrows represent findings supported by the present 16S rRNA analysis; dashed arrows represent established biological interpretations. Host immune signalling was not directly measured in this study.

While alterations in the gut microbiota have been widely reported in IBD, understanding their potential role in intestinal immune dysregulation requires consideration of both microbial composition and functional potential. Taxonomic variation between IBD and healthy individuals can provide information about changes in microbial communities, but these differences may not completely explain how microbial alterations could be related to host immune processes. Therefore, integrating 16S rRNA-based taxonomic profiling with predicted functional analysis may provide additional insights into microbial pathways that may be relevant to host–microbiota interactions and immune-related processes in IBD. This type of approach can help investigate the potential functional links between disease-associated microbiome alterations and intestinal immune responses. However, predicted functional profiles should be interpreted cautiously because they do not directly measure microbial activity [7].

Therefore, this study aimed to characterize and compare the gut microbiota profiles of individuals with IBD and healthy controls using 16S rRNA gene amplicon sequencing. This study further aimed to analyze the predicted functional potential of the microbial communities using PICRUSt2 and to identify microbial pathways associated with disease-related biological processes and potential microbiota–immune interactions in IBD. Extended analyses were conducted to prioritize candidate microbial biomarkers by integrating differential abundance results with machine-learning feature importance, evaluate a combined microbial signature, examine biomarker–pathway networks, and assess selected findings in an independent dataset. The study also examined predicted fermentation- and acetate-related pathways. As these functional profiles were predicted from 16S rRNA data, they were interpreted as functional potential rather than direct measurements of microbial activity or SCFA levels.

## 2 Materials & Methods

### 2.1 Dataset Collection

We used a publicly available database, National Center for Biotechnology Information (NCBI) Sequence Read Archive (SRA), and retrieved the raw FASTQ sequences using the SRA Toolkit [26]. The dataset contains 16S rRNA amplicon sequencing data comprising a total of 22 human stool samples, of which 11 were IBD samples and 11 were healthy control samples. The accession IDs (SRR33757848–SRR33757858) corresponded to samples collected in Shanghai, China, while (SRR37842383–SRR37842393) corresponded to samples collected in Singapore; both sets belong to gut microbiome study data. Further information including SRA accession numbers, disease status, sequencing technique, and sampling location was obtained from the SRA database and organized into a metadata sheet (Supplementary Table S1). An overview of the bioinformatics workflow used in this study is presented in Figure 2.

**Figure 2.**
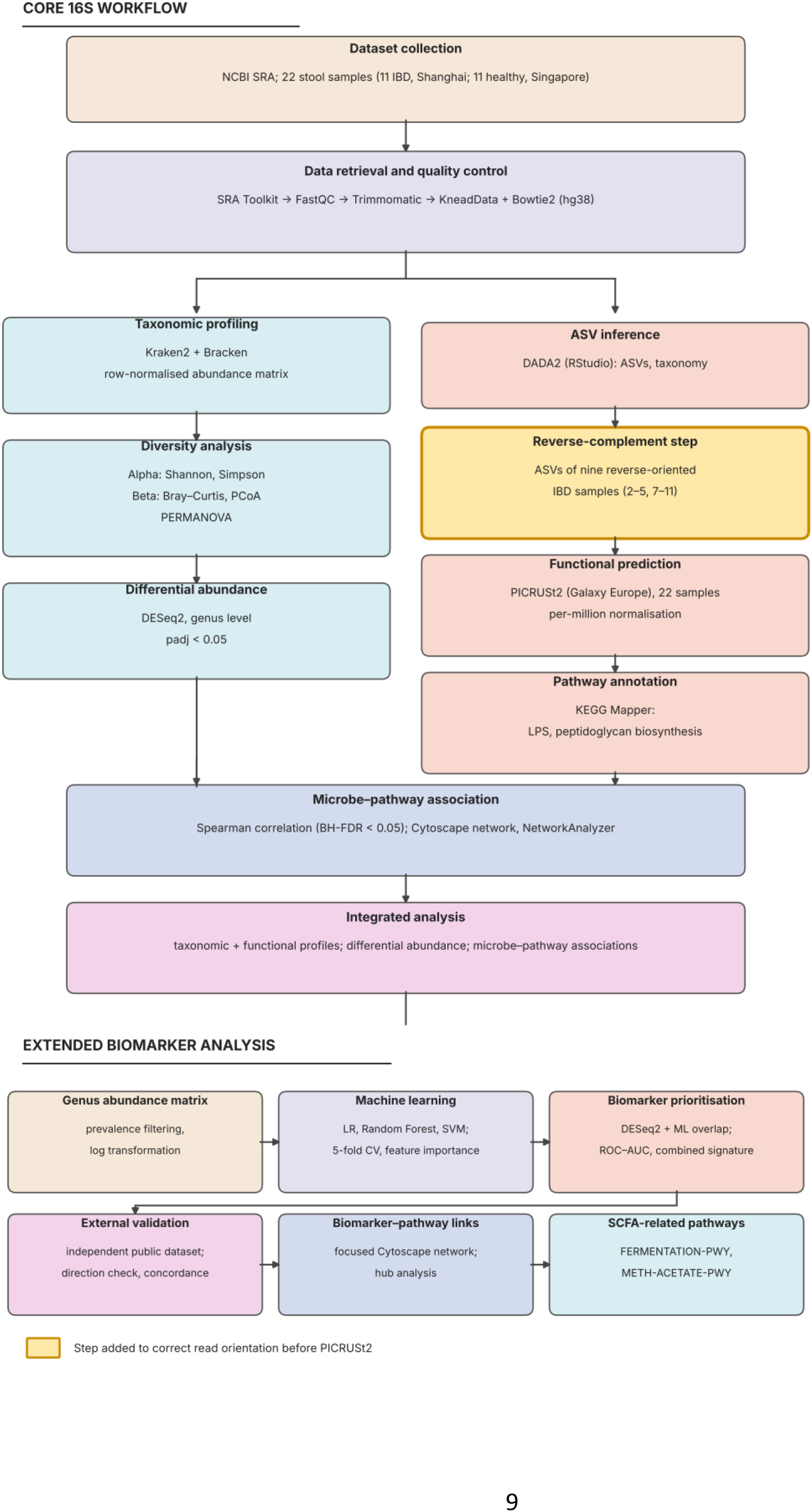

**Figure 3.**
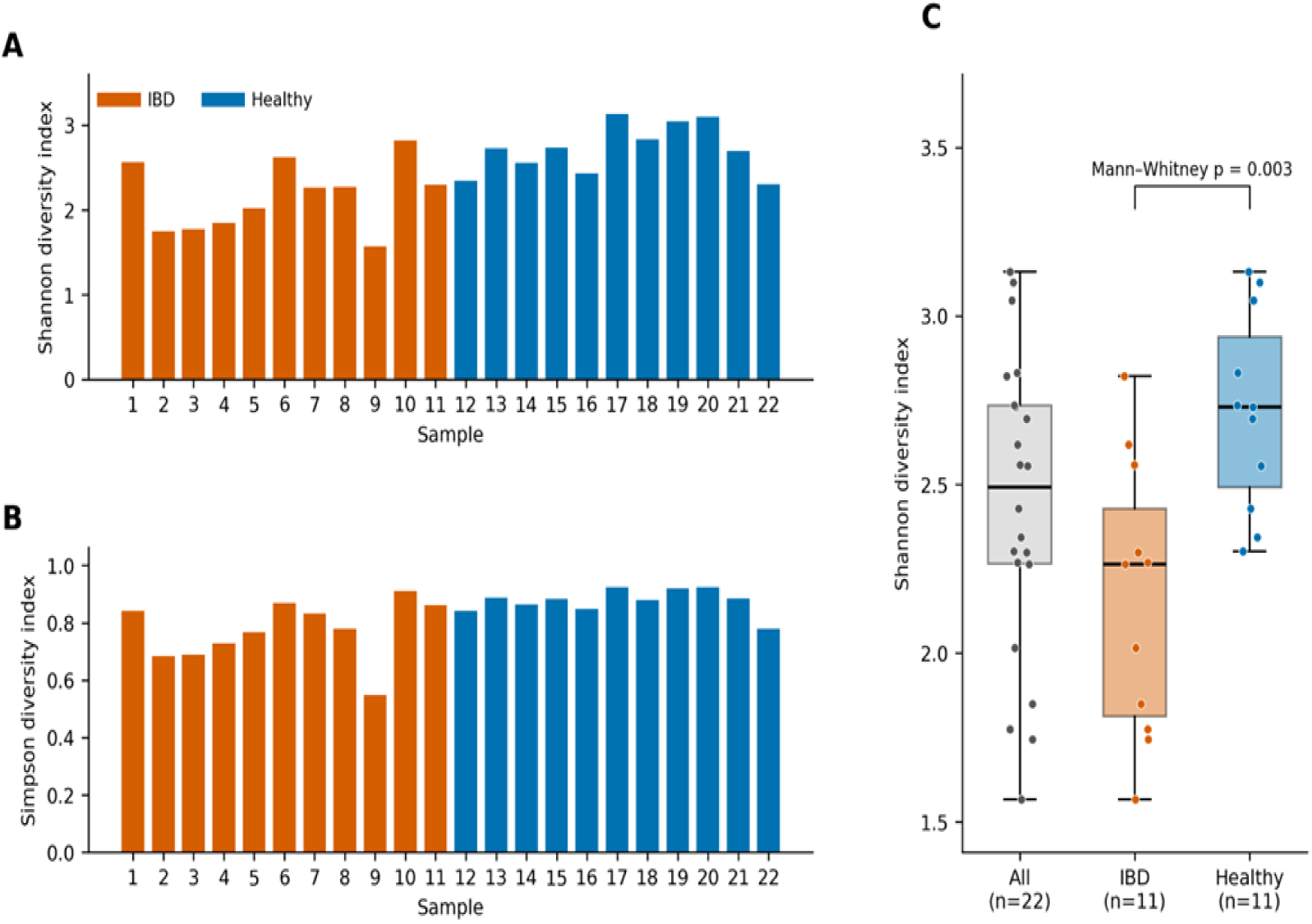
Alpha diversity of gut microbiome samples. **(A)** Shannon diversity index of the 22 gut microbiome samples (orange, IBD; blue, healthy controls). **(B)** Simpson diversity index of the 22 gut microbiome samples (orange, IBD; blue, healthy controls). **(C)** Box plots of the Shannon diversity index for all samples and separately for the IBD and healthy groups (two-sided Mann-Whitney U test, p = 0.003).

### 2.2 Computational Tools and Software

After successfully retrieving the raw sequencing data, quality assessment was performed using FastQC (v0.11.9) [27]. Adapter sequences and low-quality reads were removed using Trimmomatic (v0.39) [28], followed by host DNA removal using KneadData with Bowtie2 (v2.5.1) [29] against the human hg38 reference genome. Following quality assessment and host DNA removal, taxonomic profiling of the microbiome data was carried out: taxonomic classification was performed using Kraken2 [30], and abundance estimation was performed using Bracken [31], both implemented via Python code in a Google Colab notebook. Diversity analysis was also performed in Google Colab using the Python libraries pandas, matplotlib, seaborn, NumPy and scikit-bio (skbio.diversity), with alpha diversity assessed using the Shannon and Simpson indices and beta diversity evaluated using principal coordinates analysis (PCoA). High-resolution PCoA figures were subsequently generated in RStudio using the phyloseq and vegan packages.

For functional prediction, Amplicon Sequence Variants (ASVs) were obtained in RStudio using DADA2 [32] and used as input for PICRUSt2 [7], with functional profiling carried out via the Galaxy Europe platform [33]. KEGG Mapper [34] was subsequently used to map selected microbial pathways relevant to host–microbiota interactions. To assess associations between microbial taxa and functional features, Spearman correlation analysis was performed in RStudio, and network construction and analysis were performed in Cytoscape (v3.10.3) [35]. Differential abundance analysis was performed using the DESeq2 [36] R package, alongside the phyloseq, vegan, and ggplot2 packages, to identify taxa that differed between the IBD and healthy control groups. Finally, taxonomic and functional profiles were integrated in RStudio to support biological interpretation.

For the further extended biomarker discovery analysis, the genus-level abundance matrix was processed in Python for machine learning-based classification. Logistic Regression, Random Forest, and Support Vector Machine (SVM) classifiers were applied to distinguish IBD samples from healthy controls. Because of the limited sample size (n = 22), stratified five-fold cross-validation was implemented to maintain the class distribution across validation folds and minimize the dependence on a single train–test split. Model performance was examined using accuracy, precision, recall, F1-score, and receiver operating characteristic-area under the curve (ROC-AUC). Random Forest feature importance was estimated to rank genera according to their contribution to classification. Machine-learning-derived feature rankings were then integrated with differential abundance results to identify and prioritize key candidate microbial biomarkers. Individual candidate taxa were subsequently assessed using ROC-AUC analysis, and a combined microbial signature was evaluated to measure its classification performance. An overview of the bioinformatics workflow used in this study is presented in Figure 2.

### 2.3 Quality Control and Preprocessing

The raw FASTQ reads fetched from NCBI using the SRA Toolkit were subjected to quality control and preprocessing before downstream analysis. To assess sequence quality, FastQC (v0.11.9) was used to assess sequence quality scores, GC content, adapter content, and overrepresented sequences for each sample. Adapter sequences and low-quality reads were subsequently removed using Trimmomatic (v0.39), which was used to trim low-quality bases, adapter sequences, and unreliable positions while retaining high-quality reads. After that, host DNA removal was performed using KneadData with Bowtie2 (v2.5.1), which works by aligning reads against the human reference genome (hg38); reads that mapped to the human genome were removed. Through this process, host DNA was eliminated, leaving only high-quality, non-host microbial reads for downstream analysis.

### 2.4 Taxonomic Profiling and Diversity Analysis

For taxonomic profiling, Kraken2 (v2.1.3) was used, which is based on k-mer taxonomic classification by comparison with a reference database; it assigns reads to microbial taxa and classifies them across multiple taxonomic ranks, including phylum, class, order, family, genus, and species. For microbial abundance estimation, Bracken was used to refine the Kraken2-based output and improve abundance estimation. Bracken outputs were then merged into an abundance matrix, followed by row-wise total-sum normalization. The resulting normalized abundance matrix was used for diversity analysis (Supplementary Table S2). Diversity indices and Bray–Curtis distances were calculated from species-level Bracken abundance estimates (fraction of total reads) for all 22 samples. Alpha diversity was assessed using the Shannon and Simpson indices to evaluate diversity within samples. Differences in alpha diversity between groups were assessed using the two-sided Mann–Whitney U test. Beta diversity was assessed using the Bray–Curtis dissimilarity matrix, from which Principal Coordinates Analysis (PCoA) was generated to compare community composition between IBD and healthy control groups. Statistical significance of differences in microbial community structure between groups was assessed using Permutational Multivariate Analysis of Variance (PERMANOVA). This analysis was performed in Google Colab using Python, with the libraries pandas, matplotlib, seaborn, NumPy, and scikit-bio (skbio.diversity). PCoA coordinates were subsequently imported into RStudio for higher-resolution graphical representation, using ggplot2, and the resulting data were used for further downstream analysis.

### 2.5 Functional Profiling and Pathway Analysis

For functional profiling, Amplicon Sequence Variants (ASVs) were first generated in RStudio using DADA2, which produced representative ASV sequences, an ASV abundance table, and a taxonomy table (Supplementary Table S3). Taxonomy was assigned to the ASVs with the naive Bayesian classifier of DADA2 (assignTaxonomy) against the SILVA v138.2 training set, with both strands of each sequence tested so that reverse-oriented ASVs were also classified; a genus was assigned to 61% of the ASVs. The ASV sequences and abundance table were used as input for PICRUSt2, while the taxonomy table was retained for subsequent analysis. Because the ASV sequences of nine IBD samples (samples 2–5 and 7–11) were in reverse-complement orientation relative to the other samples, these sequences were reverse-complemented before PICRUSt2 analysis, and PICRUSt2 was then run on all 22 samples together. Predicted pathway abundances were normalised to per-million values within each sample to account for differences in sequencing depth (Supplementary Table S8b). Functional profiles were then predicted using PICRUSt2 through the Galaxy Europe platform [33]. PICRUSt2 uses phylogenetic placement of representative ASVs and reference genome information to predict gene family abundances and functional profiles, including predicted KEGG Orthologs (KOs), EC numbers, MetaCyc metabolic pathways, and pathway abundances. Predicted KOs were subsequently annotated using KEGG Mapper to identify pathways potentially associated with immune-related processes in IBD, from which selected immune-associated pathways were identified for further analysis.

### 2.6 Immune–Microbiota Association Analysis

#### Differential Abundance and Statistical Correlation analysis

To identify genera that differed between the IBD and healthy control groups, differential abundance analysis was performed using DESeq2 in RStudio. The input was the genus-level count matrix (ASV counts aggregated by genus using the taxonomy in Supplementary Table S3, excluding ASVs without a genus assignment and genera with fewer than 10 reads in total) together with the sample metadata; size factors were estimated with the “poscounts” method because of the many zero counts. DESeq2 works by using a negative binomial model to account for sequencing depth differences and biological variability. On the basis of log₂ fold change and Benjamini–Hochberg [37] adjusted p-values (false discovery rate, FDR), differentially abundant genera were determined; genera with padj < 0.05 were considered statistically significant. The differential abundance results were compiled in tabular format, and a volcano plot was generated to visualize significantly different genera between IBD and healthy control groups.

To find the relationship between microbial genera and immune-related pathways, Spearman’s rank correlation analysis was performed in RStudio by combining the genus abundance matrix with the PICRUSt2-derived pathway abundance table. Genus abundance was expressed as relative abundance (ASV counts aggregated at genus level using the taxonomy in Supplementary Table S3), pathway abundances were used as per-million values, and genera detected in at least two samples were included. For each microbe–pathway pair, the Spearman correlation coefficient (ρ) was calculated to assess the strength and direction of their association. Correlation p-values were corrected using the Benjamini–Hochberg false discovery rate (FDR) method to reduce the potential for false-positive associations. Microbe–pathway associations with an FDR-adjusted p-value < 0.05 were considered statistically significant. The resulting correlation matrix was used for network construction and further analysis.

### 2.7 Network Construction and Integration

For better understanding and clear visualization of the associations between bacterial genera and predicted functional pathways, Cytoscape (v3.10.3) was used. This was done by exporting the significant correlations (FDR-adjusted p < 0.05) from the Spearman analysis as an edge list, which was then imported into Cytoscape for network analysis. Bacterial genera and functional pathways were represented as nodes, and the edges indicated statistically significant associations between them. The network facilitated the identification of strongly connected genera and major pathway hubs, simplifying the complex microbe–pathway associations. Finally, to support biological interpretation, taxonomic and functional information were combined in RStudio. This integrated dataset included taxonomic information (genus, phylum, family), PICRUSt2-predicted immune-related pathways, DESeq2 statistics (log₂FC and adjusted p-values), and significant Spearman correlation results (ρ and FDR). This integrated framework provided a basis for assessing microbial composition alongside predicted functional potential in IBD. The microbe–pathway network tables were analysed in Cytoscape (NetworkAnalyzer) [38] to calculate node degree and centrality measures, and the network and other figures were generated in Python (matplotlib) from the supplementary tables.

#### 2.8.1 Preparation of Genus-Level Abundance Matrix

To perform the extended analysis, first, a genus-level abundance matrix was prepared from the existing taxonomic abundance data. In the matrix, rows represented the samples, and columns represented bacterial genera with their respective abundance values. For classification, the information from the Sample and Group columns was retained, and the dataset included 22 samples, comprising 11 IBD and 11 healthy controls.

#### 2.8.2 Filtering of Low-Abundance and Low-Prevalence Genera

After that, the genus-level abundance matrix was used for the further filtering step. The filtering step was performed to remove low-abundance or low-prevalence genera before downstream analysis. A prevalence threshold of ≥20% of samples and a relative abundance threshold of ≥0.01% were applied. Those genera that did not meet the criteria for minimum prevalence and abundance were subsequently excluded from the ML analysis. Therefore, filtering was performed before the downstream analysis.

#### 2.8.3 Data Normalization and Feature Scaling

For ML preprocessing, filtered genus-level abundance data were utilized, and the abundance data were represented in relative abundance format. After that, log transformation was applied, e.g., X =np.log1p(X). In the log-transformed microbial features, feature scaling/standardisation was applied where required by the model. Then, the output of this was an ML-ready feature matrix, and the preprocessing was performed within the cross-validation framework where appropriate to avoid data leakage.

#### 2.8.4 Definition of the Prediction Problem

The prediction problem was defined in the form of binary classification: Healthy = 0 and IBD = 1. X represented the microbial genus features, and y represented the disease status/group. In the ML models, X was used in the form of predictors/features, and y was used as the target variable. For example: X = df.drop(columns=[“Group”]) and y = df[“Group”].

#### 2.8.5 Cross-Validation Strategy

In the dataset, the number of samples was 22, with 11 IBD and 11 Healthy controls. Because of the limited sample size, the conventional 80:20 train–test split was not used as a primary approach. Instead, stratified five-fold cross-validation was used. The purpose of stratification was to maintain the class distribution of the IBD and Healthy groups in each fold. Hence, this approach was applied to the limited dataset for model performance evaluation.

#### 2.8.6 Train Logistic Regression

Logistic Regression was trained as a baseline model, and the genus abundance features were used as input. The model generated the predicted probability of IBD classification. Model performance was evaluated using the following metrics: Accuracy, Precision, Recall, F1-score, and ROC-AUC. This evaluation was performed within the framework of stratified five-fold cross-validation.

#### 2.8.7 Random Forest Classification

Now, the Random Forest was applied as another ML classifier, and the classification task was IBD vs Healthy controls. The same genus-level microbial features were used as input. Random Forest was evaluated using the stratified five-fold cross-validation framework. Then, the feature importance was calculated from the Random Forest model. Feature importance was used to rank the bacterial genera on the basis of their contribution to the classification, and the ranked genera were later utilized for candidate biomarker prioritization.

#### 2.8.8 Model Comparison and Candidate Biomarker Prioritization

For IBD vs Healthy classification, multiple machine-learning classifiers were used, and the models considered in this were Logistic Regression (LR), Random Forest (RF), and Support Vector Machine (SVM). This was done to compare the classification performance of different ML models. For model classification, the same filtered and preprocessed genus-level microbial abundance features were used. Model performance was evaluated by Accuracy, Precision, Recall, F1-score, and ROC-AUC, and the evaluation was done using the stratified five-fold cross-validation framework. Because the dataset was small, with only 22 samples, small differences in performance between models should not be overinterpreted.

On the basis of feature importance and ranking from the ML analysis, important bacterial genera were identified. After that, these ML-derived results were integrated with the DESeq2 differential abundance results. The purpose of this integration was to identify the bacterial genera that showed both statistical differential abundance evidence and ML predictive importance. Then, on the basis of this combined evidence, the genera were prioritized in the form of candidate microbial biomarkers.

#### 2.8.9 ROC-AUC Analysis and Integrated Biomarker Selection

ROC-AUC analysis was used for further evaluation of prioritized candidate microbial genera, and ROC analysis was performed to analyze the ability of individual candidate taxa to distinguish IBD samples from Healthy controls. Microbial abundance was utilized to generate the ROC curve and calculate the area under the curve (AUC) for individual taxa. For classification performance, sensitivity and specificity were also considered. A combined microbial signature was also estimated using selected candidate taxa to distinguish IBD from Healthy controls. The combined microbial signature was evaluated using ROC-AUC and related classification performance measures.

The results of DESeq2, ML feature importance, and ROC-AUC analysis were integrated, and important/candidate microbial genera from different evidence layers were compared. On the basis of the combined evidence, candidate taxa were prioritized. Taxa that were consistently important across multiple analytical approaches were selected for the final candidate microbial biomarker panel. The integrated analysis combined differential abundance evidence from DESeq2, predictive importance from ML, and classification performance from ROC-AUC. Then, the final selected taxa were used for subsequent biomarker–pathway integration.

#### 2.8.10 Biomarker–Pathway Integration

The final selected candidate microbial biomarkers were integrated with significant PICRUSt2-predicted functional pathways. The purpose was to assess the relationship between candidate biomarker taxa and predicted microbial functions and then compare and link the selected biomarker taxa with significant microbial pathways identified from the functional analysis. Using the available correlation results, microbe–pathway associations were examined, and Spearman correlation was used to evaluate associations between microbial taxa and functional pathways. Multiple-testing correction was applied using FDR, and significant associations were identified based on the predefined significance/FDR criterion. This analysis focused particularly on pathways relevant to the study, including LPS biosynthesis and peptidoglycan biosynthesis. A biomarker–pathway integration table was prepared containing candidate biomarker, direction of abundance change, associated pathway, correlation coefficient (rho), p-value, and FDR. Hence, these associations were subsequently used for network construction and biological interpretation.

#### 2.8.11 Biomarker-Focused Network Construction and Hub Analysis

For biomarker-focused network construction, already available microbe–pathway correlation results were utilized. In the extended analysis, candidate microbial biomarkers were included in the network. The network was mainly focused on LPS biosynthesis and peptidoglycan biosynthesis. In the network, microbial taxa and pathways were represented in the form of nodes, and significant microbe–pathway correlations were represented in the form of edges. Candidate biomarkers were separately distinguished in the network. The network was constructed and visualized in Cytoscape v3.10.3.

After that, the constructed network was further analyzed in Cytoscape Network Analyzer, and three network parameters, degree, betweenness centrality, and closeness centrality, were calculated to evaluate the connectivity and importance of the nodes within the network. These parameters were used to identify the highly connected/important nodes.

#### 2.8.12 External Validation of Candidate Microbial Biomarkers

For external validation, an independent publicly available 16S rRNA dataset was used, and the dataset represented an IBD cohort. The samples in the dataset were stool samples. The cohort included Crohn’s disease (CD) and control/healthy samples, with a total of 43 samples (23 Crohn’s disease and 20 controls; MicrobiomeAnalyst example IBD dataset, file ibd_data.zip, https://www.microbiomeanalyst.ca/MicrobiomeAnalyst/resources/data/ibd_data.zip). The dataset was used for external validation of candidate microbial biomarkers identified by the discovery analysis. In the independent dataset, the candidate genera/taxa from the discovery analysis were evaluated. Candidate taxa were first checked for availability in the validation dataset, and the taxa available in the external validation dataset were subjected to between-group abundance comparison between the CD and control groups. The comparison was performed using the Mann–Whitney U test for abundance comparison, and FDR correction was applied. The direction of abundance change was also compared between the discovery and validation datasets. Therefore, this analysis was intended to assess whether the identified microbial associations showed independent external support.

#### 2.8.13 Biomarker Concordance Analysis

The candidate biomarker findings from the discovery and independent validation datasets were compared to assess their concordance. The direction of abundance change of the available candidate taxa was compared between the discovery dataset and the validation dataset. The discovery analysis compared IBD and healthy controls, whereas the external validation analysis compared CD and control groups. Candidate taxa showing the same direction of abundance change across both datasets were considered concordant. This analysis was used to assess whether the findings identified in the discovery analysis showed consistent independent support in the external validation dataset.

#### 2.8.14 SCFA- related pathway analysis

For SCFA-related pathway analysis, the per-million-normalised predicted pathway abundance data obtained from PICRUSt2 were used as input, with a focus on fermentation- and acetate-related pathways. Specifically, FERMENTATION-PWY and METH-ACETATE-PWY were analyzed. The predicted pathway abundances were compared between the IBD and Healthy groups using the Mann–Whitney U test. Group-wise pathway abundance was summarized using median values, and the direction of abundance change between the two groups was determined. FDR correction was applied to the obtained P-values to account for multiple comparisons, and statistical significance was assessed based on an FDR threshold of < 0.05. The group-wise distribution of the predicted pathway abundances was visualized using boxplots with individual sample values.

## 3 Results

### 3.1 Taxonomic Analysis of the Gut Microbiota

#### Alpha Diversity Analysis

Shannon and Simpson indices from the alpha diversity analysis of gut microbiome samples indicated that the Shannon index (Figure 3A) varied across all 22 samples, and its value range goes from the lowest value of 1.57 in Sample 9 to the highest value of approximately 3.13 in Sample 17. Most samples ranging from 2.0 to 2.8 were more clustered within this range. The Simpson index (Figure 3B) also showed the same trend as the Shannon index. The values ranged from 0.55 to 0.93. Sample 9 showed low diversity, while Samples 10, 17, 19, and 20 showed the highest diversity scores. The Shannon diversity box plot (Figure 3C) revealed the distribution of Shannon diversity values across all samples. The horizontal line inside the box represents the median Shannon score, which lies at approximately 2.49, while the interquartile range falls between 2.27 and 2.73. The values ranged from 1.57 (Sample 9) to 3.13 (Sample 17); the box plots are shown for all samples and separately for the IBD and healthy groups. Alpha diversity was significantly lower in the IBD group than in the healthy control group (Shannon index: median 2.27 vs 2.73, Mann–Whitney U = 15, p = 0.003; Simpson index: median 0.78 vs 0.88, U = 17, p = 0.005; 11 IBD and 11 healthy samples). The complete alpha diversity results are provided in Supplementary Table S4.

#### Beta Diversity Analysis

To observe the variation in microbial community composition between the healthy control and IBD groups, beta diversity analysis was performed. The Bray–Curtis distance matrix was used, and Principal Coordinates Analysis (PCoA) was performed to visualize the beta diversity pattern. Figure 4 showed that the IBD and healthy control samples were separated mainly along PC1 (33.6% of the variance; PC2 = 23.4%), with the IBD samples at lower and the healthy samples at higher PC1 values (Mann–Whitney p = 0.003). Three IBD samples (6, 7, and 11), however, were located close to the healthy samples (Figure 4B). The PERMANOVA analysis (Pseudo-F = 6.125, p < 0.001; 9999 permutations) indicated that the microbial community composition differed significantly between the groups. However, because the IBD and healthy samples originated from different cohorts, this difference may partly reflect cohort and sequencing differences rather than disease status alone. The Bray–Curtis distance matrix, PCoA coordinates, and complete PERMANOVA results are provided in Supplementary Tables S5, S6, and S7, respectively.

**Figure 4.**
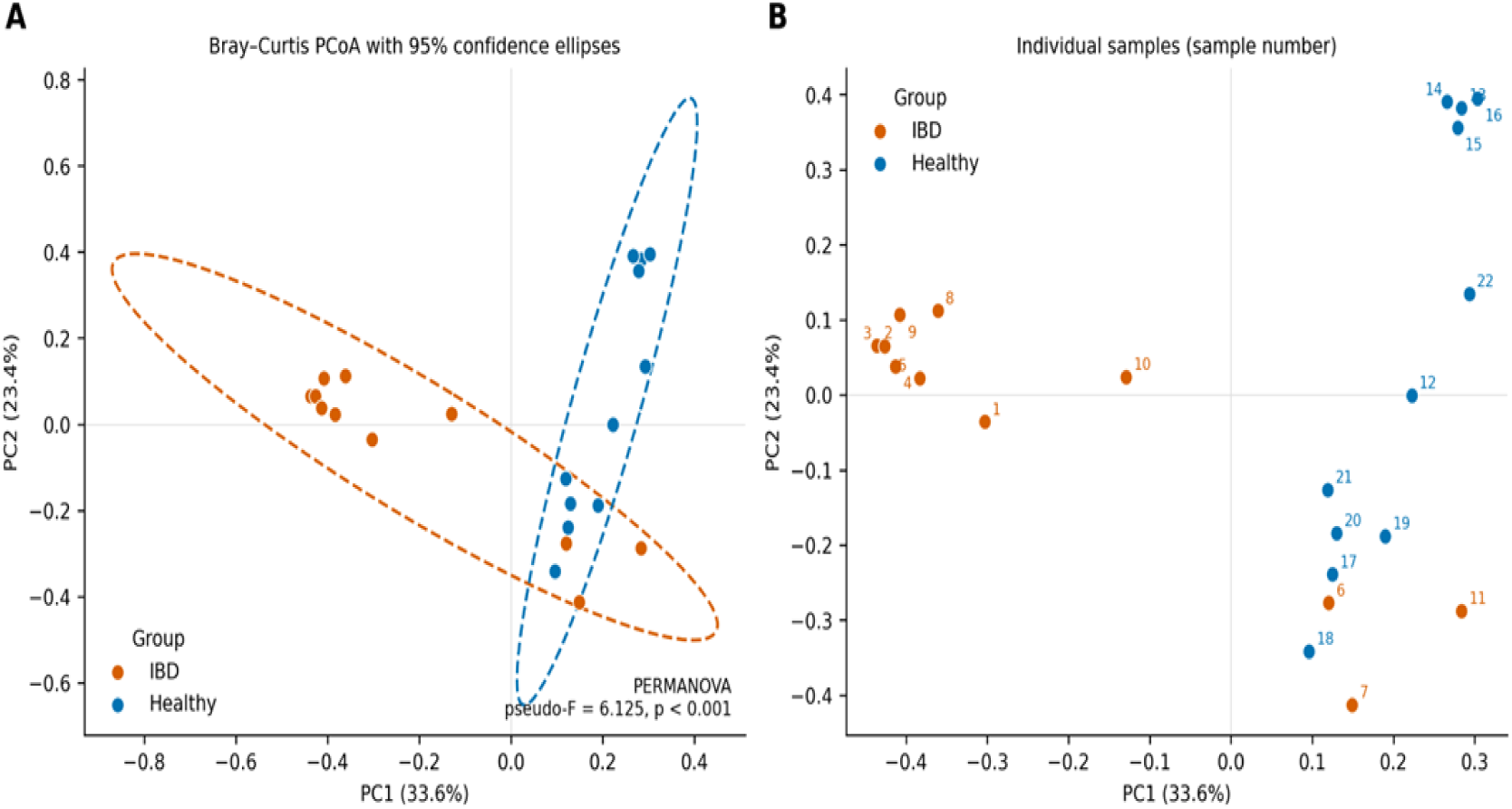
Beta diversity of the gut microbiota based on Bray-Curtis dissimilarity. **(A)** Principal coordinates analysis (PCoA) showing the IBD (orange) and healthy control (blue) samples, with dashed lines indicating the 95% confidence ellipse of each group. **(B)** PCoA showing the individual samples, labelled by sample number. PERMANOVA indicated that microbial community composition differed significantly between the groups (pseudo-F = 6.125, p < 0.001; 9999 permutations). PCI and PC2 explain 33.6% and 23.4% of the variance, respectively.

### 3.2 Functional Profiling

Based on the functional prediction performed by PICRUSt2, the functional profiling of the gut microbiome samples predicted the functional potential of the microbial communities across all samples. PICRUSt2 generated the predicted KEGG Orthologs (KOs) and the pathway abundance matrix for all samples. After reverse-complementing the reverse-oriented ASVs, pathway predictions were obtained for all 22 samples (282–354 predicted pathways per sample; 392 pathways in total). The predicted KO IDs were utilized in KEGG Mapper to map them to multiple metabolic pathways. The complete PICRUSt2 pathway abundance data are provided in Supplementary Table S8, and the per-million-normalised values used for all downstream comparisons are provided in Supplementary Table S8b. From the list of predicted pathways, two immune-associated pathways were filtered: Lipopolysaccharide biosynthesis and Peptidoglycan biosynthesis (Table 1; Supplementary Table S9). Both pathways were also present in the PICRUSt2 pathway abundance table and were selected for further analysis.

**Table 1.**
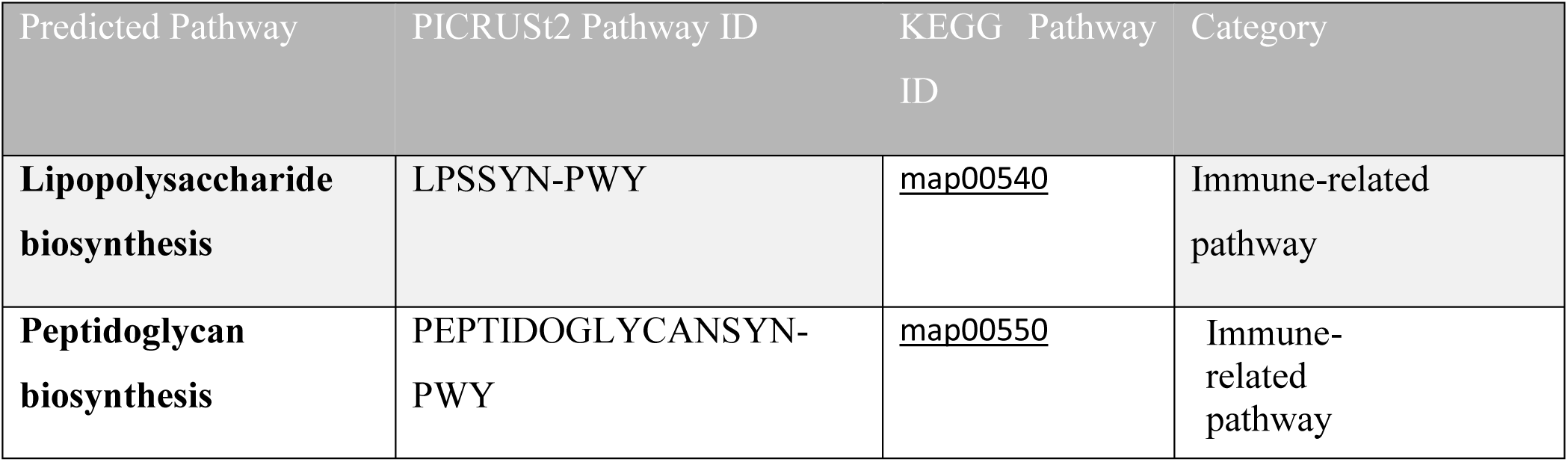
Immune-related pathways identified from PICRUSt2-predicted KEGG Orthologs through KEGG Mapper annotation.

### 3.3 Immune–Microbiota Association Analysis

#### Differential Abundance Analysis

To distinguish the bacterial genera between the IBD and healthy control groups, differential abundance analysis was performed using DESeq2. Of the 186 genera tested by DESeq2, 44 were significantly differentially abundant (FDR-adjusted p < 0.05); 32 were lower and 12 were higher in the IBD group. Most of these genera (27 of 44) were detected in only one of the two groups, which produces very large log₂ fold changes and very small adjusted p-values; these results therefore mainly reflect the presence or absence of genera in one group rather than graded abundance differences, and only 17 of the significant genera were detected in both groups. Ten representative genera (those with the smallest adjusted p-values among genera detected in at least five samples) are shown in Table 2: eight showed negative log₂ fold-change values (lower in IBD) and two showed positive values (higher in IBD). The significant DESeq2 genera are provided in Supplementary Table S10, while the complete DESeq2 results are provided in Supplementary Table S11. The volcano plot (Figure 5A) visualizes the distribution of differentially abundant genera according to log₂ fold change and −log₁₀(FDR).

**Table 2.**
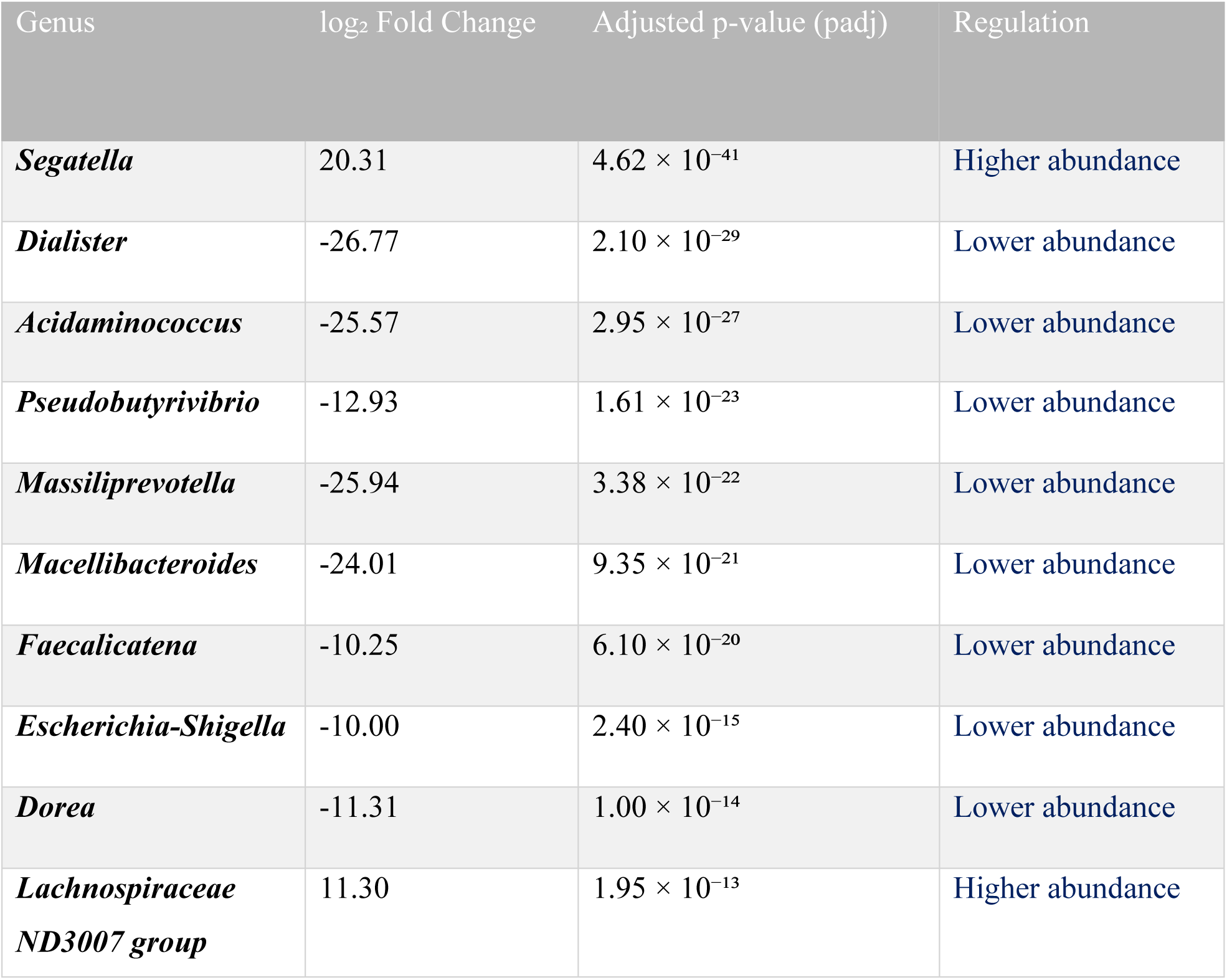
Top differentially abundant bacterial genera identified between disease and healthy groups using DESeq2.

**Figure 5.**
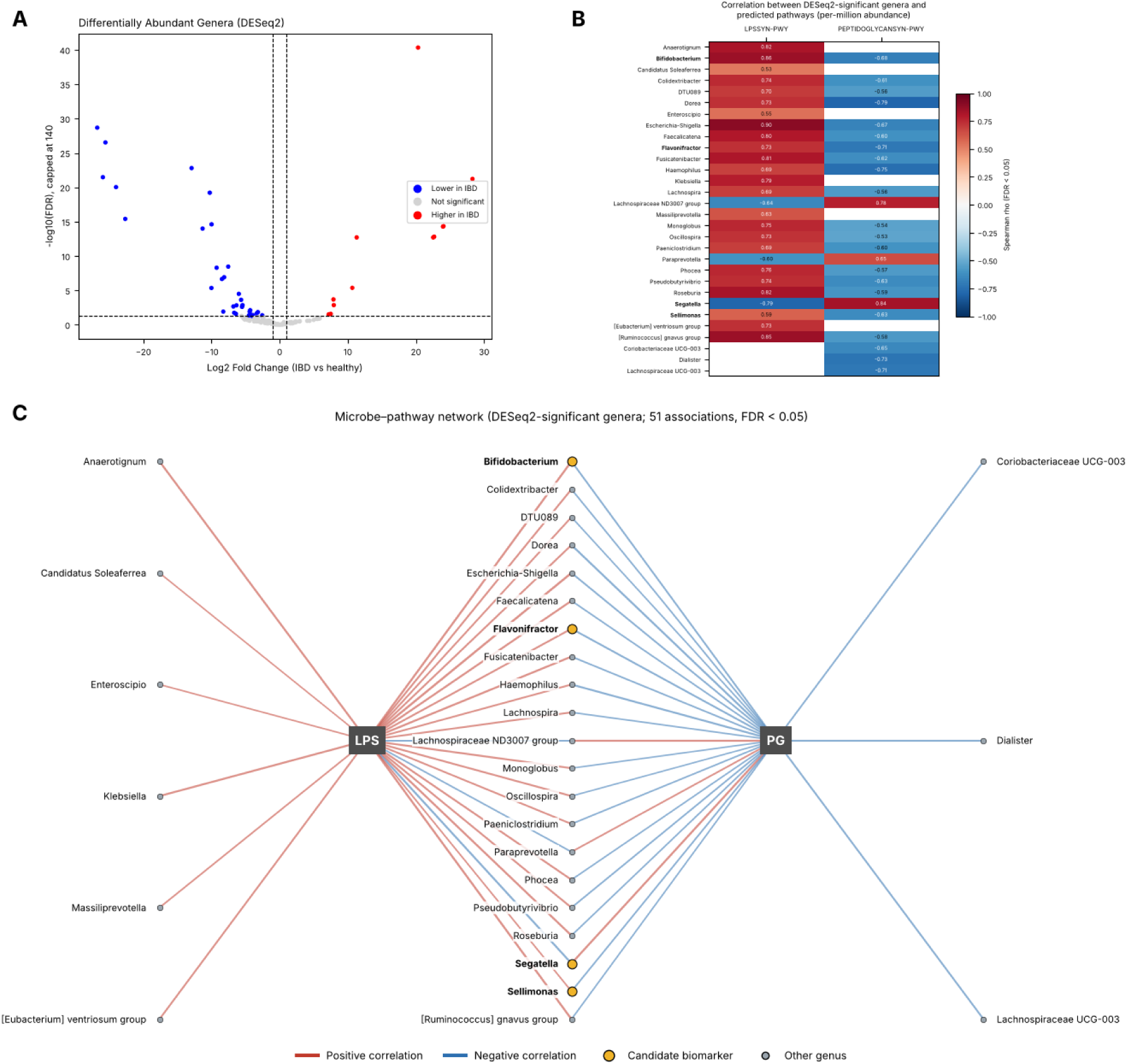
Differential abundance and microbiota-immune pathway associations. (A) DESeq2 volcano plot of differentially abundant bacterial genera between IBD and healthy controls (genera absent from one group have very large fold changes and should be read as presence/absence differences). (B) Heatmap of FDR-significant Spearman correlations between DESeq2-signifìcant genera and the predicted LPSSYN-PWY and PEPTIDOGLYCANSYN-PWY pathways (per-million-normalised pathway abundance); candidate biomarkers are in bold, (C) Network of the 51 significant microbe-pathway associations; red edges, positive correlations; blue edges, negative correlations; candidate biomarkers are highlighted.

### 3.4 Correlation and Network Analysis

Spearman rank-order correlation analysis was performed between genus relative abundances and per-million-normalised pathway abundances, and this analysis showed the associations between microbial genera and LPSSYN-PWY and PEPTIDOGLYCANSYN-PWY. A total of 84 significant correlations were identified, with the significance criterion of FDR-adjusted p < 0.05: 47 with LPSSYN-PWY and 37 with PEPTIDOGLYCANSYN-PWY. Of the 47 correlations with LPSSYN-PWY, 44 were positive (ρ = 0.5095 to 0.9004; mean ρ = 0.6799) and three were negative (Segatella, ρ = −0.7933; Lachnospiraceae ND3007 group, ρ = −0.6406; Paraprevotella, ρ = −0.5983). Of the 37 correlations with PEPTIDOGLYCANSYN-PWY, 33 were negative (ρ = −0.8171 to −0.5248; mean ρ = −0.6316) and four were positive (Segatella, ρ = 0.8357; Lachnospiraceae ND3007 group, ρ = 0.7773; UCG-003, ρ = 0.6649; Paraprevotella, ρ = 0.6539). The strongest positive correlations with LPSSYN-PWY were observed for Escherichia-Shigella (ρ = 0.9004), Bifidobacterium (ρ = 0.8561), and the [Ruminococcus] gnavus group (ρ = 0.8536), and the strongest negative correlations with PEPTIDOGLYCANSYN-PWY for Bacteroides (ρ = −0.8171), Dorea (ρ = −0.7930), and Haemophilus (ρ = −0.7520). Because the correlated genera were mostly lower in abundance in the IBD group, LPSSYN-PWY was detected in only one of the 11 IBD samples (compared with all 11 healthy samples), and peptidoglycan biosynthesis was relatively higher in the IBD group (median 8,524 vs 7,333 per million; Mann–Whitney p = 0.0013), these correlations mainly reflect differences between the two groups; the few genera with the opposite pattern (notably Segatella, which was present in all IBD samples and absent from the healthy samples) are those that were higher in the IBD group. Figure 5B showed the relationship between the DESeq2-significant genera and the two predicted pathways, LPSSYN-PWY (Lipopolysaccharide biosynthesis) and PEPTIDOGLYCANSYN-PWY (Peptidoglycan biosynthesis). The significant microbe–pathway correlations are provided in Supplementary Table S12.

Cytoscape network analysis was used to visualize the significant microbe–pathway associations. Figure 5C represents the two pathway nodes in the network, LPSSYN-PWY and PEPTIDOGLYCANSYN-PWY, which act as central hubs. Multiple microbial genera are connected to these pathway nodes, and 21 genera were connected to both pathways.

### 3.5 Integrated Taxonomic and Functional Analysis

To assess the association between bacterial genera and selected predicted immune-related pathways, the taxonomic and functional results were combined. This integrated analysis included Genus, Phylum, Family, log₂FoldChange, padj, Pathway, Rho, and FDR. log₂FoldChange and padj represented by the DESeq2 results, while the functional results were represented by the predicted pathways. The correlation results were represented by Spearman ρ (Rho) and FDR. Lipopolysaccharide biosynthesis (LPSSYN-PWY) and Peptidoglycan biosynthesis (PEPTIDOGLYCANSYN-PWY) were the selected pathways. A number of genera were associated with these pathways. Some representative genera in the integrated dataset, including Bifidobacterium, Colidextribacter, Dorea, Escherichia-Shigella, Flavonifractor, Haemophilus, Roseburia, and the [Ruminococcus] gnavus group, are presented in Table 3, and the complete integrated dataset is provided as Supplementary Table S14. All genera shown were lower in abundance in the IBD group; their correlations were positive with lipopolysaccharide biosynthesis and negative with peptidoglycan biosynthesis.

**Table 3.** Representative differentially abundant bacterial genera associated with selected predicted immune-related pathways.

| Genus | Phylum | Log<sub>2</sub><br>Fold<br>Change | Adjusted *p*-<br>value (*padj*) | Predicted pathway | Spearman<br>$\rho$ |
| --- | --- | --- | --- | --- | --- |
| **Bifidobacterium** | Actinomycetota | −4.315 | $2.93 \times 10^{-2}$ | Lipopolysaccharide biosynthesis | 0.856 |
| **Colidextribacter** | Bacillota | −4.331 | $7.56 \times 10^{-3}$ | Lipopolysaccharide biosynthesis | 0.745 |
| **Dorea** | Bacillota | −11.315 | $1.00 \times 10^{-14}$ | Peptidoglycan biosynthesis | −0.793 |
| **Escherichia-Shigella** | Pseudomonadota | −9.995 | $2.40 \times 10^{-15}$ | Lipopolysaccharide biosynthesis | 0.900 |
| **Flavonifractor** | Bacillota | −5.660 | $2.35 \times 10^{-4}$ | Lipopolysaccharide biosynthesis | 0.733 |
| **Haemophilus** | Pseudomonadota | −6.338 | $1.45 \times 10^{-3}$ | Peptidoglycan biosynthesis | −0.752 |
| **Roseburia** | Bacillota | −5.518 | $2.44 \times 10^{-3}$ | Lipopolysaccharide biosynthesis | 0.825 |
| **[Ruminococcus] gnavus group** | Bacillota | −7.549 | $3.41 \times 10^{-9}$ | Lipopolysaccharide biosynthesis | 0.854 |

### 3.6 Machine-Learning Classification and Feature Importance

Machine-learning models were analysed using stratified five-fold cross-validation. Logistic Regression attained an accuracy of 0.864, precision of 1.000, recall of 0.727, an F1-score of 0.842, and a ROC–AUC of 0.917. Random Forest attained scores of 1.000 for all five metrics in each fold. SVM achieved an accuracy of 0.773, precision of 1.000, recall of 0.545, an F1-score of 0.706, and a ROC–AUC of 0.901 (Table 4; Supplementary Tables S16 and S18). The highest mean Random Forest feature-importance scores were observed for *Sodalis* (0.0956), *Enterococcus* (0.0879), *Segatella* (0.0738), *Sellimonas* (0.0687), and *Adlercreutzia* (0.0531) (Supplementary Table S19).

**Table 4.** Cross-validation performance of machine-learning classifiers for distinguishing IBD from healthy controls.

| Model | Accuracy | Precision | Recall | F1-score | ROC–AUC |
| --- | --- | --- | --- | --- | --- |
| Logistic Regression | **0.864** | **1.000** | **0.727** | **0.842** | **0.917** |
| Random Forest | **1.000** | **1.000** | **1.000** | **1.000** | **1.000** |
| SVM | **0.773** | **1.000** | **0.545** | **0.706** | **0.901** |

Note: Random Forest achieved a score of 1.000 for each metric in all five cross-validation folds. Fold-wise results are provided in Supplementary Table S18.

### 3.7 Candidate Biomarker Prioritization

Five genera were recognized in the original analysis as both DESeq2-significant and important in the machine-learning analysis: Sellimonas, Bifidobacterium, Flavonifractor, Segatella, and Paludicola. After the genus taxonomy was corrected, four of them (Segatella, Sellimonas, Flavonifractor, and Bifidobacterium) remained DESeq2-significant, whereas Paludicola did not (padj = 0.188). On the basis of combined evidence, Segatella, Sellimonas, Bifidobacterium, and Flavonifractor were classified as high-priority candidates, whereas Paludicola, which was selected by the machine-learning and ROC analyses (Random Forest rank 6; AUC 0.984) but was not DESeq2-significant, was classified as low priority (Supplementary Table S20). The integrated biomarker panel ranked Segatella first, followed by Sellimonas, Paludicola, Bifidobacterium, and Flavonifractor (Supplementary Table S24; Figure 6 [c]).

**Figure 6.**
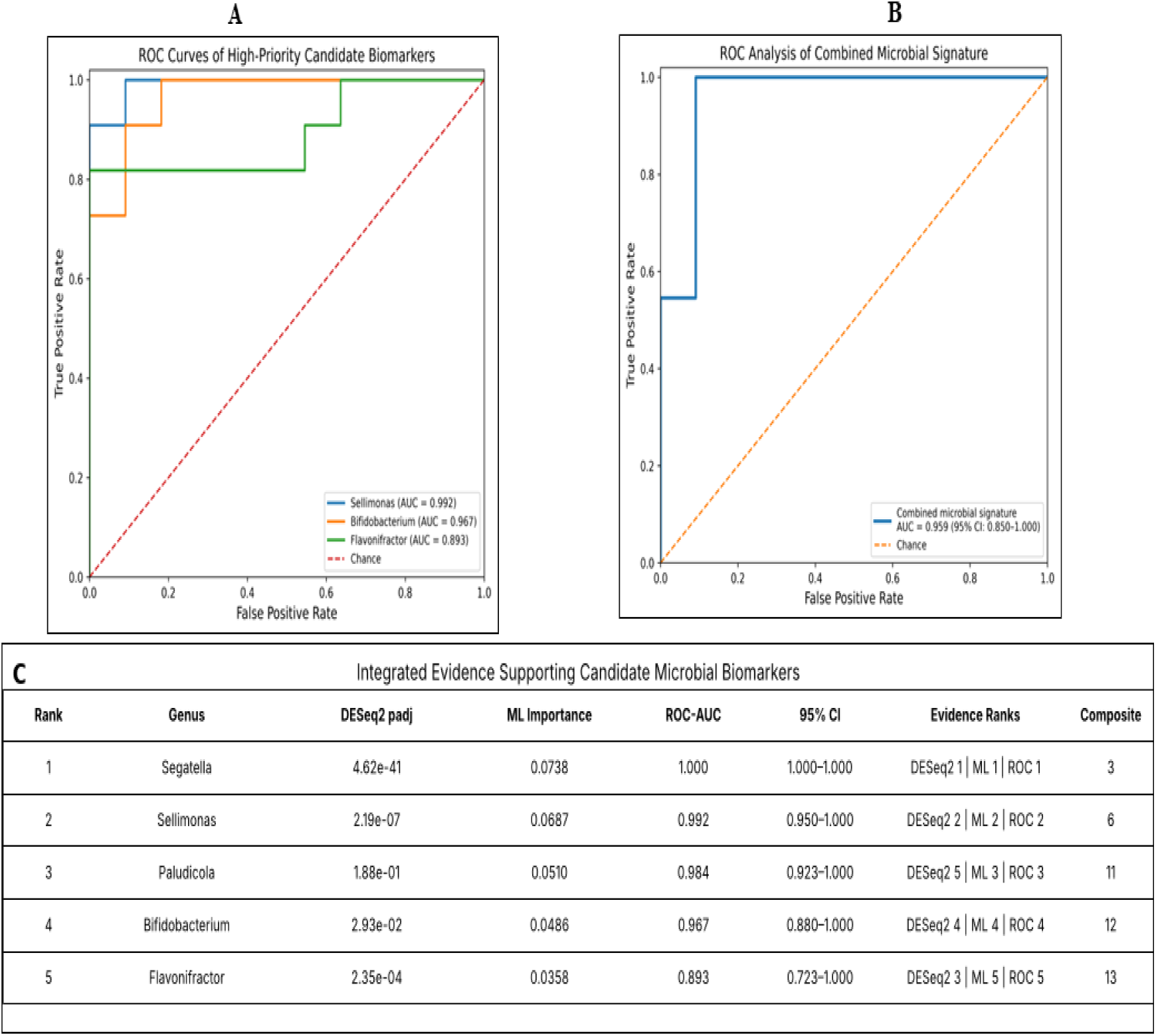
ROC analysis and integrated evidence for candidate microbial biomarkers. **(A)** ROC curves for high priority candidate biomarkers. **(B)** ROC curve for the combined microbial signature. **(C)** Integrated evidence used to rank candidate biomarkers.

### 3.8 ROC–AUC Analysis and Combined Microbial Signature

Among the high-priority candidate biomarkers, Segatella separated the groups perfectly (ROC–AUC = 1.000; it was present in all IBD samples and absent from all healthy samples), and among the other candidates Sellimonas demonstrated the ROC–AUC of 0.9917, followed by Bifidobacterium (0.9669) and Flavonifractor (0.8926) (Figure 6[A]; Supplementary Table S21). The combined signature of these three genera (Sellimonas, Bifidobacterium, and Flavonifractor, the high-priority candidates in the initial ranking) achieved an ROC–AUC of 0.9587 (95% CI: 0.85–1.00), with an accuracy of 0.9545, sensitivity of 1.000, specificity of 0.9091, and F1-score of 0.9565 (Figure 6[B]; Supplementary Tables S22 and S23).

### 3.9 Biomarker-Focused Network and Hub Analysis

The biomarker-focused network contained 51 significant microbe–pathway associations among 30 DESeq2-significant genera, including 27 associations with LPS biosynthesis (24 positive, 3 negative) and 24 with peptidoglycan biosynthesis (21 negative, 3 positive). Among the five candidate biomarkers, Bifidobacterium, Flavonifractor, and Sellimonas were each connected to both pathways (network degree = 2), with positive correlations with LPS biosynthesis (ρ = 0.856, 0.733, and 0.589, respectively) and negative correlations with peptidoglycan biosynthesis (ρ = −0.685, −0.714, and −0.632). Segatella was also connected to both pathways but with the opposite pattern (ρ = −0.793 with LPS biosynthesis and ρ = 0.836 with peptidoglycan biosynthesis), whereas Paludicola had no significant pathway associations. LPS biosynthesis had the highest network degree (27), followed by peptidoglycan biosynthesis (24) (Supplementary Tables S25–S28) (Figure 7A, B)

**Figure 7.**
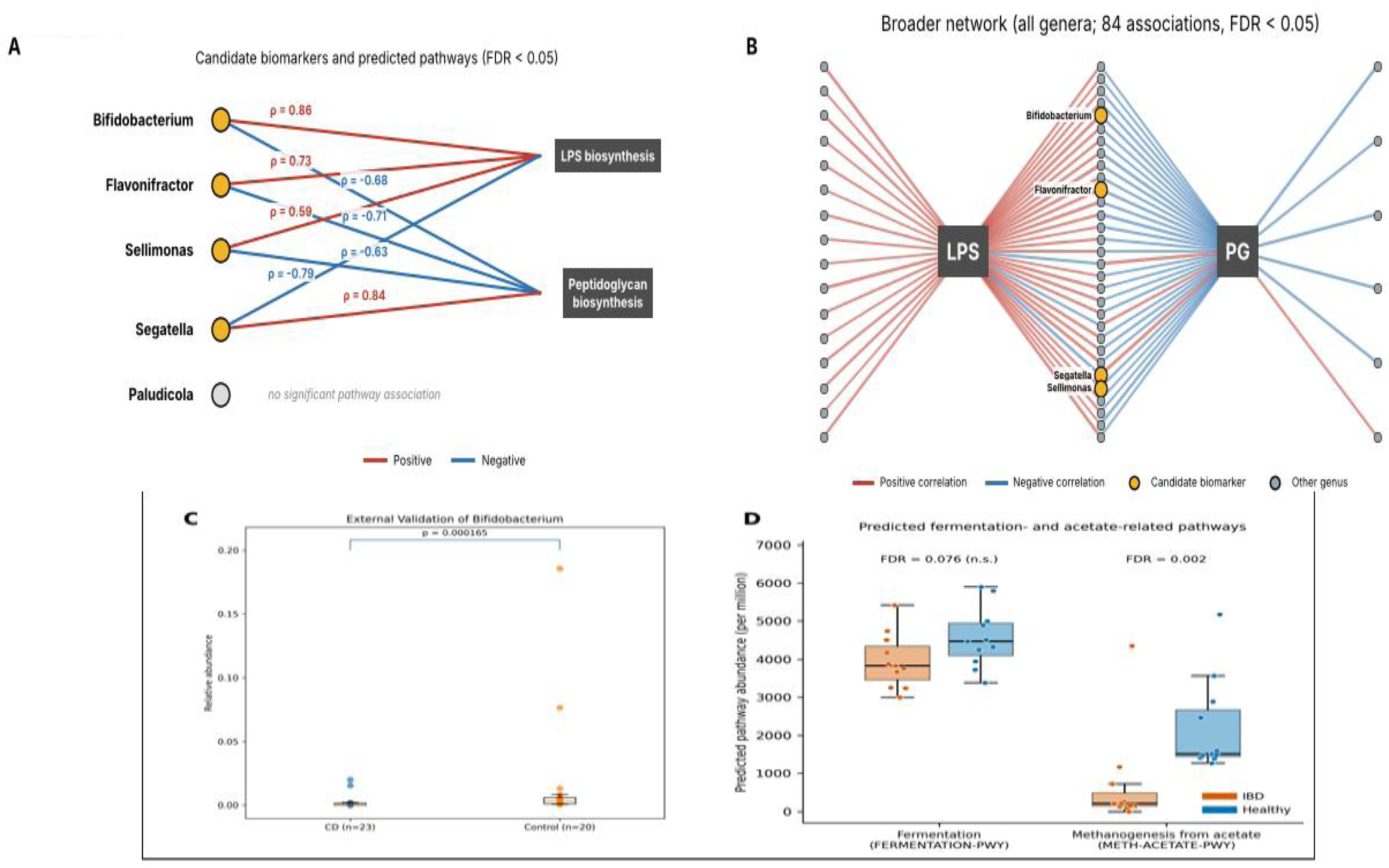
Extended biomarkor analyses. (A) Associations of the candidate biomarkers with the predicted lipopolysaccharide (LPS) and peptidoglycan biosynthesis pathways (Speannan p shown; FDR < 0.05). Bifidobacterium, Flavonifractor and Sellimonas were positively correlated with LPS and negatively correlated with peptidoglycan biosynthesis, Segatella showed the opposite pattern, and Paludicola showed no significant pathway association. (B) Bro der network of all 84 significant microbe-pathway associations (FDR < 0.05); red edges indicate positive and blue edges negative correlations, and candidate biomarkers are highlighted in gold. (C) External validation of Bifidobacterium in Crohn’s disease ICD, n = 23) and control (n = 20) samples (Mann-Whitney U test, p = 0.000165). (D) Predicted fermentation- and acetate-related (methanogenesis from acetate) pathway abundances, expressed per million, in the IBD and healthy groups; FDR-adjusted p-values are shown (FERMENTATION-FWY, FDR= 0.076; METH-ACETATE-PWY, FDR = 0.002).

### 3.10 External Validation and Biomarker Concordance

External validation was performed using an independent dataset comprising 23 patients with Crohn’s disease (CD) and 20 healthy controls. *Bifidobacterium* abundance was significantly higher in the control group than in the CD group (Mann–Whitney U = 75, p = 0.000165; Figure 7C; Supplementary Table S29). Of the five candidate biomarkers assessed, *Bifidobacterium* was the only genus detected in the external dataset, and its direction of change was concordant with the discovery analysis. Thus, one of the five candidate biomarkers could be evaluated and showed concordant abundance differences in the independent dataset (Supplementary Tables S30–S31).

### 3.11 SCFA-Related Predicted Pathway Analysis

After normalising pathway abundances to per-million values, the predicted acetate-related pathway (METH-ACETATE-PWY, methanogenesis from acetate) was lower in the IBD group than in healthy controls (median 205.93 vs 1,505.07 per million; FDR-adjusted p = 2.05 × 10⁻³), whereas the predicted fermentation pathway (FERMENTATION-PWY) did not differ significantly between the groups (median 3,824.58 vs 4,470.74 per million; FDR-adjusted p = 0.076; Figure 7D; Supplementary Table S32).

## Discussion

This current study examined the association between gut microbiota composition and predicted immune-related functional pathways in inflammatory bowel disease (IBD) through bioinformatics analysis of 16S rRNA sequencing data. An integrated workflow including microbial diversity, taxonomic profiling, differential abundance analysis, PICRUSt2-based functional prediction, KEGG pathway mapping, Spearman correlation analysis, and Cytoscape network construction was used. Altogether, the analysis identified variations in microbial diversity, differentially abundant bacterial genera, and predicted immune-associated pathways, providing a basis for exploring potential microbiota–immune interactions and microbial dysbiosis in IBD.

The alpha diversity analysis demonstrated variation in microbial diversity across the gut microbiome samples, indicating heterogeneity among the microbial communities examined. In the present dataset, alpha diversity was significantly lower in the IBD group than in healthy controls, consistent with previous observations of altered microbial diversity in IBD, although this comparison is confounded by cohort of origin (see Limitations). A systematic meta-analysis of 13 IBD microbiome studies reported lower alpha diversity in both Crohn’s disease and ulcerative colitis compared with non-IBD controls, supporting altered microbial diversity as a consistent feature of IBD-associated dysbiosis [10]. Large-scale microbiome studies have likewise documented substantial alterations in the gut microbial ecosystem, community structure, and functional characteristics in IBD [11,12]. Overall, the variation observed in the present study is broadly aligned with previous reports of altered microbial diversity in IBD, although the magnitude and direction of these changes may differ across cohorts. Differences in disease phenotype, treatment exposure, sample characteristics and other sources of individual-level variation may contribute to this variability [10,11]. In beta diversity analysis, the IBD and healthy control samples were separated mainly along the first principal coordinate in the Bray–Curtis PCoA, indicating differences in microbial community composition between the groups, although substantial inter-individual variation was also present. Comparable heterogeneity has been reported in IBD studies, where differences in microbial community composition between IBD and healthy individuals may coexist with substantial inter-individual variation within the disease group [13]. Ryan et al. reported differences in colonic microbiota composition associated with IBD and inflammation, while also highlighting heterogeneity in host–microbiota interactions [13]. In larger cross-cohort assessments, significant differences in Bray–Curtis beta diversity between IBD and healthy groups have been reported; however, cohort-related factors were also found to significantly influence microbial community composition [14]. In the present study, the PERMANOVA result was statistically significant (Pseudo-F = 6.125, p < 0.001), indicating that community composition differed between the IBD and healthy control groups; however, because the two groups originated from different cohorts, this separation may partly reflect cohort-related and sequencing differences rather than disease status alone.

The functional profiling in the current study identified lipopolysaccharide biosynthesis and peptidoglycan biosynthesis within the predicted microbial pathways selected for immune - related analysis. These pathways are of particular interest in IBD because bacterial structural components can interact with the host innate immune system and play a role in inflammatory signalling. Lipopolysaccharide from Gram-negative bacteria can participate in pattern - recognition receptor-mediated inflammatory responses, whereas bacterial peptidoglycan is recognized by innate immune receptors and can influence intestinal immune responses[5,8,9].Earlier investigations have highlighted the close interaction between the gut microbiota and host immune system in IBD, indicating that changes in microbial components and their recognition may contribute to intestinal inflammation [5].Hence, the prediction of pathways related to lipopolysaccharide and peptidoglycan biosynthesis in the current study might indicate a potential functional link between the gut microbiota and immune-related processes in IBD. However, as these pathways were inferred from 16S rRNA sequencing using PICRUSt2[7], they should be regarded as predicted functional potential rather than direct evidence of pathway activity or increased production of these bacterial components.

The differential abundance patterns noted in the current study showed similarities and differences when compared with earlier studies of IBD-associated gut dysbiosis. Although Coprococcus showed higher abundance in the current study, previous findings have reported alterations in the abundance of this genus in IBD, including reduced abundance in crohn’s disease [15]. Likewise, alterations in the abundance of Dialister, Sutterella, and other bacterial genera have been reported across different IBD cohorts, although the pattern of these changes has varied between studies [15,16]. Such differences in the abundance patterns of particular genera might reflect the heterogeneous nature of the gut microbiome in IBD. Differences in disease subtype, disease activity, medication exposure, and other factors might contribute to variations between cohorts. Hence, the differentially abundant genera identified in the current study can be considered features of microbial dysbiosis within the studied cohort rather than universal microbial markers of IBD.

The associations observed here between bacterial genera and the predicted lipopolysaccharide (positive) and peptidoglycan (negative) biosynthesis pathways should be interpreted in light of earlier studies connecting microbial functional characteristics with immune-related processes in IBD, although the direction of genus-level associations has differed between studies. In a study of mucosa-associated microbiota in Crohn’s disease, functional inference from 16S rRNA sequencing showed that predicted LPS biosynthesis pathways were increased in patients before treatment, and these pathways correlated negatively with genera such as Roseburia, Ruminococcus, and Lachnospira [17]. Emerging metatranscriptomic evidence has also shown increased expression of LPS biosynthesis functions within microbes associated with Crohn’s disease, along with pro-inflammatory host responses, further supporting the association of microbial LPS-related functions with intestinal inflammation [18]. Peptidoglycan-derived bacterial components are likewise recognized by innate immune receptors, such as NOD1, NOD2, and TLR2, and can impact inflammatory signaling in the intestinal environment [8,9]. Hence, the microbe–pathway associations observed in the present study may reflect a correlation between variation in bacterial community composition and the predicted functional potential of immune-relevant bacterial pathways. Yet, these associations should not be interpreted as evidence that the correlated genera directly induce LPS or peptidoglycan production, especially because the functional profiles in the current study were estimated from 16S rRNA sequencing using PICRUSt2 [7]. Notably, all genera listed in Table 3 were lower in abundance in the IBD group, and LPSSYN-PWY was detected in only one of the 11 IBD samples; the correlations therefore largely reflect differences between the two groups (including differences in cohort and sequencing characteristics) rather than an enrichment of LPS- or peptidoglycan-producing taxa in IBD. The relatively higher peptidoglycan biosynthesis in the IBD group after depth normalisation, which produced the negative correlations with these genera, should likewise be interpreted cautiously.

The integrated taxonomic and functional analysis gave a broader perspective on the relationship between microbial abundance patterns and predicted microbial functions in IBD.

Existing studies on multi-omics have shown that taxonomic and functional alterations in the gut microbiome can take place together during IBD activity, along with integrated analyses identifying microbial and functional properties associated with disease-related dysregulation [11]. Likewise, cross-cohort multi-omics analysis identified interlinked bacterial, functional, and metabolite properties associated with IBD, supporting the value of integrated taxonomic and functional information while assessing microbial dysbiosis [14]. More recently, large-scale metagenomic analysis across multiple IBD cohorts has additionally shown that IBD-associated microbial populations are not uniform and that their functional characteristics can differ according to species and strain variants [19]. In the current study, the integration of differential abundance and microbe–pathway correlation results thus gave a broader view of the relationship between altered bacterial genera and predicted immune-related microbial functions. Still, these associations should be interpreted cautiously because functional profiles were estimated via 16S rRNA sequencing using PICRUSt2 [7], and the integrated associations identified in the present cohort should not be considered universal microbial–functional markers of IBD.

In the extended biomarker analyses, differential abundance results were combined with machine-learning feature importance to identify candidate genera. Sellimonas, Bifidobacterium, Flavonifractor, Segatella, and Paludicola were initially identified by both approaches; after correction of the genus taxonomy, Paludicola was no longer DESeq2-significant (padj = 0.188), although it remained among the top Random Forest features, while the integrated evidence ranking still placed Segatella first. Random Forest achieved perfect scores across the five cross-validation folds; however, this result should be interpreted cautiously because the discovery dataset was small (n = 22), and machine-learning models developed with small datasets can show optimistic performance and require validation in larger, independent cohorts [20,21]. The combined three-genus signature also demonstrated high classification performance in this dataset, but this internal result alone does not establish clinical diagnostic value [22]. Because the signature genera were prioritized using the full discovery dataset (DESeq2 and Random Forest importance) before the combined model was evaluated, the reported ROC–AUC of 0.959 is likely optimistic and should be regarded as exploratory.

The biomarker-focused network analysis showed significant associations between four of the five candidate genera and the predicted LPS and peptidoglycan biosynthesis pathways: Bifidobacterium, Flavonifractor, and Sellimonas were positively associated with LPS and negatively associated with peptidoglycan biosynthesis, whereas Segatella showed the opposite pattern. However, not all prioritized candidate genera had significant pathway associations in the network (Paludicola had none). These network results represent correlations with predicted functional pathways and should not be interpreted as evidence of direct microbial activity or causation [7,8,9].

The external dataset showed that *Bifidobacterium* abundance was significantly higher in healthy controls than in patients with Crohn’s disease. This direction was concordant with the discovery analysis; however, only one of the five candidate genera could be evaluated in the external dataset. Hence, this finding provides limited validation of the candidate panel and should be interpreted cautiously. Differences in cohort characteristics, disease subtype, sample size, and taxonomic detection may influence the reproducibility of microbiome findings across datasets [23,24]. Further validation in larger, independent cohorts is needed.

After normalisation for sequencing depth, the predicted acetate-related pathway (methanogenesis from acetate, METH-ACETATE-PWY) was lower in the IBD group than in healthy controls, whereas the predicted fermentation pathway did not differ significantly between the groups. Because METH-ACETATE-PWY represents methanogenesis from acetate, a pathway of methanogenic archaea that consumes acetate, its lower predicted abundance should not be interpreted as reduced acetate production; these findings may indicate differences in the predicted microbial functional potential between the groups. However, these pathways were inferred from 16S rRNA data using PICRUSt2 and do not represent direct measurements of short-chain fatty acids or pathway activity [7]. SCFAs are produced through microbial fermentation, and changes in SCFA levels have been reported in IBD [25]. Therefore, the predicted pathway differences in this study require confirmation using direct metabolomic or shotgun metagenomic analyses.

The current study has certain limitations that should be considered when interpreting the findings. The moderately small sample size of 22 samples might limit the statistical power and generalizability of the observed microbial patterns. In addition, disease status was confounded with cohort of origin: all IBD samples came from one study (Shanghai, China) and all healthy controls from another (Singapore), and information on age, sex, and treatment was not included in the metadata used. Moreover, the sequences of the IBD and healthy samples did not overlap: none of the 2,266 ASVs was detected in both groups, and nine IBD samples contained reads in the opposite orientation that had to be reverse-complemented before PICRUSt2 analysis, indicating differences in sequencing and read processing between the cohorts. Differences between the groups, including the perfect Random Forest classification and the near-absence of LPSSYN-PWY in the IBD samples, may therefore partly reflect technical, batch, or population differences rather than disease biology. In addition, only 61% of the ASVs could be classified to genus level, so the genus-level results describe the classified fraction of each community, and 27 of the 44 DESeq2-significant genera were detected in only one of the two groups, so their fold changes and p-values mainly reflect the presence or absence of a genus in a cohort (which, given the cohort confounding and non-overlapping ASVs, may be technical in origin). The machine-learning analysis used Kraken2/Bracken genus abundances, whereas differential abundance testing and the microbe–pathway correlations used ASV-based genus counts, and the two were not fully concordant for all genera. The perfect Random Forest scores across the five cross-validation folds should also be interpreted cautiously, as models developed with small datasets can show optimistic performance and require validation in larger, independent cohorts [20,21]. Moreover, the functional profiles were predicted from 16S rRNA sequencing using PICRUSt2 and thus represent predicted functional potential rather than directly measured microbial activity or SCFA levels [7]. The observed microbe–pathway correlations do not establish causal relationships between specific bacterial genera and predicted functional pathways. External validation was limited because only one of the five candidate genera could be evaluated, although its abundance direction was concordant with the discovery analysis. In addition, the external cohort compared Crohn’s disease with controls rather than IBD overall. Differences in cohort characteristics, disease phenotype, treatment exposure, and taxonomic detection might affect the reproducibility of microbiome findings across datasets [23,24]. Hence, the microbial and functional associations identified in the current study require validation in larger independent cohorts and, where feasible, using shotgun metagenomic, metatranscriptomic, or metabolomic approaches. Finally, two genus-level abundance tables were used: Kraken2/Bracken estimates for diversity analysis, and DADA2 ASV-derived genus counts for differential abundance and correlation analyses. Genus-level estimates and genus names differed between the two approaches, so results based on different tables are not directly comparable.

## Conclusion

The current work gave an integrated overview of gut microbial dysbiosis and predicted immune-related functional potential in inflammatory bowel disease (IBD). This assessment identified differences in microbial diversity and the abundance of specific bacterial genera between IBD and healthy controls, together with predicted lipopolysaccharide and peptidoglycan biosynthesis pathways. Correlation and network analyses identified associations between altered bacterial genera and the predicted lipopolysaccharide (positive) and peptidoglycan (negative) biosynthesis pathways, which largely reflect differences between the groups and should be interpreted cautiously. Extended analyses combined differential abundance with machine-learning feature importance to prioritize candidate genera and evaluated a three-genus microbial signature. External validation was limited to *Bifidobacterium*, whose abundance direction was concordant with that in the discovery analysis. Of the two SCFA-related pathways examined, only the predicted acetate-related pathway (methanogenesis from acetate) was lower in the IBD group. Through combining the taxonomic, functional, network-level, and extended biomarker analyses, this work contributes to an in-depth knowledge of potential microbiota–immune interactions in IBD and offers a hypothesis-proposing framework for identifying microbial and functional features associated with IBD-associated dysbiosis. Despite the fact that these functional associations are based on PICRUSt2 predictions, they do not prove direct microbial activity or causation, and the pathway results do not represent direct measurements of SCFAs. In summary, these findings contribute as a basis for further research and confirmation of microbial–functional associations in larger independent cohorts.

## Data availability

The raw 16S rRNA amplicon sequencing data analysed in this study are publicly available in the NCBI Sequence Read Archive (SRA) under accession numbers SRR33757848–SRR33757858 (IBD samples; Shanghai, China) and SRR37842383–SRR37842393 (healthy control samples; Singapore). Sample-level information is provided in Supplementary Table S1. The independent validation dataset (43 stool samples: 23 Crohn’s disease and 20 controls) was obtained from the MicrobiomeAnalyst example IBD dataset (ibd_data.zip; https://www.microbiomeanalyst.ca/MicrobiomeAnalyst/resources/data/ibd_data.zip, accessed 2026). All supplementary tables (S1–S32, including S8b and S27b) accompany this manuscript.

## Code availability

The analysis code (Python notebooks and R scripts), supplementary tables, and Cytoscape network files and session are available at https://github.com/amanzehra18-oss/ibd-16s-microbiome (archived at Zenodo DOI: https://doi.org/10.5281/zenodo.23359812).

## Ethics statement

This study used only publicly available, de-identified sequencing data. No new samples were collected, and no additional ethical approval was required.

## Author contributions

A.Z. conceived the study, performed the data retrieval, bioinformatic and statistical analyses, prepared the figures and wrote the manuscript. E.N. supervised the study and provided guidance on methodology and interpretation. Both authors reviewed and approved the final manuscript.

## Acknowledgements

The authors thank Biotecnika Info Labs for providing the platform and support for this work.

## Use of AI-assisted tools

AI-assisted tools were used to help write and check analysis code, verify the re-analysis, and edit the manuscript text. The authors reviewed and verified all analyses, results and text, and take full responsibility for the content of this work.

