## Supplementary Data 1 for "Integrated 16S rRNA Bioinformatics Analysis of Gut Microbiota, Predicted Pathways, and Candidate Biomarkers in Inflammatory Bowel Disease": Supplementary_Data_Index.txt

Supplementary Data Index

S2 Bracken species-level abundance for all 22 samples (genus and species matrices, relative abundance and read counts).

Tables S1-S14: Original core 16S analysis supplementary tables, retained without renumbering.

Extended analysis tables:
S15 Filtered genus abundance matrix used for machine-learning analysis.
S16 Logistic Regression, Random Forest, and SVM model comparison.
S17 Logistic Regression out-of-fold sample predictions.
S18 Random Forest five-fold cross-validation results.
S19 Random Forest feature-importance ranking.
S20 ML-important taxa and integrated evidence for candidate prioritization.
S21 Individual candidate-taxon ROC-AUC results.
S22 Combined microbial signature performance.
S23 Combined-signature out-of-fold predictions.
S24 Integrated biomarker panel.
S25 Biomarker-focused Cytoscape edge table.
S26 Biomarker-focused Cytoscape node table.
S27 Cytoscape NetworkAnalyzer metrics.
S28 Network hub analysis summary.
S29 Independent-dataset external validation results.
S30 Biomarker concordance details.
S31 Biomarker concordance summary.
S32 PICRUSt2-predicted SCFA-related pathway comparison.

Workflow Step 17 (disease-subtype analysis) was not included because it was not performed.

Version note (corrected release):
- S6: the Group column was previously inverted relative to S1; it now lists samples 1-11 as Disease (IBD) and samples 12-22 as Healthy. PCoA coordinates are unchanged.
- S10: now contains only the 49 genera with FDR-adjusted p < 0.05 (previously 59 rows, 10 of which were not significant). S11 still contains all 175 tested genera.
- S20: DESeq2_significant and Evidence_score corrected for the 10 genera that are not significant (padj >= 0.05). The five candidate biomarkers and their priorities are unchanged.
- S10 and S11: first column labelled "Genus"; S6: first column labelled "Sample".
- File names for S4 and S5 corrected (removed double full stop).


Version note (v3, functional analysis regenerated):
- S8: PICRUSt2 re-run on reorientated ASVs (1014 ASVs from nine IBD samples were reverse-complemented). All 22 samples now have predicted pathways (392 pathways). S8b: the same table normalised to per-million (per-sample total) to correct for the lower sequencing depth of the IBD samples.
- S12, S13, S14, S25-S28, S32 and the Functional_association / Evidence_score columns of S20 were recalculated from the per-million table (Spearman correlation, Benjamini-Hochberg FDR within each pathway, genera present in >= 2 samples; genus abundance from ASV counts aggregated using the S3 annotation). S13 = significant correlations of DESeq2-significant genera (padj < 0.05). S27 was computed with NetworkX, not Cytoscape NetworkAnalyzer.
- These tables are re-derivations; please confirm them by re-running the original analysis code on S8b.

Version note (v4, diversity re-run with all 22 samples):
- S4, S5, S6, S7 were recalculated for all 22 samples (11 IBD, 11 healthy) from the species-level Bracken files (fraction_total_reads; Shannon index in natural log). Sample 20 was previously omitted because its Bracken file was saved as .xlsx rather than .csv. The 21 previously reported Shannon and Simpson values were reproduced.
- S5 rows and columns are now in numeric sample order (1-22). In the previous version the matrix values were in numeric order but labelled in text-sorted order (10, 11, ..., 1, 21, 22, 2, ...), and the S6 group labels were assigned in the same wrong order; the previous PERMANOVA (pseudo-F = 1.754, p = 0.091) was therefore based on mismatched labels.
- S7: PERMANOVA with 9999 permutations (previously 999).

Version note (v5):
- S27b: Cytoscape NetworkAnalyzer node table (undirected) for the broader microbe-pathway network shown in Figure 7B (65 nodes, 95 edges; Supplementary Table S12). S27 refers to the biomarker-focused network of Figure 5C (67 edges).

Version note (v6):
- S27: replaced by the Cytoscape NetworkAnalyzer node table (undirected) of the biomarker-focused network of Figure 5C (41 nodes, 67 edges). S28 hub summary updated with these values; Segatella and Paludicola have no significant pathway association and are not part of the network.

Version note (v7, release for manuscript and bioRxiv):
- S2: replaced. It previously contained only the Bracken species reports of 21 samples as separate sheets; it now contains a combined matrix for all 22 samples (sheets: README, Samples, Genus_fraction, Genus_reads, Species_fraction, Species_reads). Genus = first word of the Bracken species name; genus relative abundance is the sum of species-level fraction_total_reads. The sample 20 report was added (Shannon 3.098, consistent with S4).

Version note (v8, genus taxonomy re-assigned; genus-level results regenerated):
- Reason: for nine IBD samples (2, 3, 4, 5, 7, 8, 9, 10, 11) the ASVs are in reverse orientation, so no genus had been assigned to them in S3 (0% genus-level assignment for these samples). Taxonomy was re-assigned with the SILVA 138.2 training set (silva_nr99_v138.2_toGenus_trainset, naive Bayesian classifier, 8-mers, 100 bootstraps, minBoot 50), trying both strands (tryRC). Genus is now assigned to 61% of ASVs (previously 39%). Of the 892 ASVs that had a genus before, 874 keep the identical genus, 18 are now unassigned and none changed genus.
- S3: replaced by the re-assigned taxonomy (2,266 ASVs).
- S10, S11: DESeq2 (R; poscounts size factors, Wald test, IBD vs healthy, genera with >= 10 reads, "Unclassified" excluded) re-run on the corrected genus table: 186 genera tested, 44 with padj < 0.05 (12 higher, 32 lower in IBD). 27 of the 44 are absent from one of the two groups, so their log2 fold changes and p-values are very large and should be read as presence/absence differences. Genera detected in a single sample are replaced by DESeq2's outlier handling and have baseMean 0 / padj NA.
- S12-S14, S20, S24-S28, S27b: recalculated from the corrected genus table (Spearman correlation with the per-million pathway table S8b, Benjamini-Hochberg FDR within each pathway, genera present in >= 2 samples). 84 significant genus-pathway correlations (S12); 51 of these involve DESeq2-significant genera (S13, S25); biomarker-focused network: 32 nodes, 51 edges (S27); broader network: 55 nodes, 84 edges (S27b).
- S24: Paludicola is no longer DESeq2-significant (padj = 0.19) and is now low priority in S20; the five-genus panel is otherwise unchanged. S15-S19, S21-S23 and S29-S31 are based on the Bracken genus abundances (S2) and are unchanged.
- S2, S4-S9 and S32 are unchanged.
